# Reticulocyte Infection Leads to Altered Behaviour, Drug Sensitivity and Host Cell Remodelling by *Plasmodium falciparum*

**DOI:** 10.1101/862169

**Authors:** Renugah Naidu, Trang TT Chu, Jaishree Tripathi, Yang Hu, Gowtham Subramanian, Jie Xin Tong, Pallavi Tripathi, Kong Fang, Kevin SW Tan, Chwee Teck Lim, Jerry K.Y. Chan, Zbynek Bozdech, Rajesh Chandramohanadas

**Author notes:** Corresponding Author: Rajesh Chandramohanadas.

## Abstract

Plasmodia are host-specific, both at the organism and cellular levels. During asexual development, *Plasmodium spp.* infect cells of erythroid lineage, with an overall propensity towards reticulocytes. This applies to even *Plasmodium (P.) falciparum*, the most common causative agent of human malaria, implications of which remain unexplored. Herein, for the first time, we characterize the developmental stages and features of *P. falciparum* cultured *in vitro* in young reticulocytes (CD71^+^) in comparison to standard normocyte (CD71^-^) cultures. We demonstrate that there are notable differences in the patterns of invasion, development and sensitivity to potent antimalarials (such as artemisinin and dihydroartemisinin) for parasites residing in CD71^+^ reticulocytes. Through a transcriptomic approach, we report that *P. falciparum* parasites are able to sense the host cell environment, and calibrate their metabolic and host cell remodelling pathways through differential gene expression. These results form an exciting avenue on which hitherto unexplored interactions between *Plasmodium spp* and different stages of host red blood cells could be investigated in the broader contexts of drug resistance, host tropism and zoonosis.

**Author Summary:** Parasites causing malaria infect red blood cells for development and proliferation during asexual development. This asexual erythrocytic stage determines higher parasite densities and eventual disease manifestation. Although the most virulent species of Plasmodium infecting humans known as *Plasmodium falciparum* is able to infect red blood cells of all ages, these parasites show a preference for younger blood cells. Of note, the biochemical and biophysical properties of young and adult red blood cells vary significantly. Herein, we undertook a comparative profiling of invasion process, parasite development and drug response of *Plasmoddium falciparum* in two host cells: young red blood cells (reticulocytes) and mature red blood cells (normocytes). We demonstrate that *P. falciparum* infects human reticulocytes with higher affinity and demonstrate differential sensitivity to drugs such as artemisinin while they reside within reticulocytes. Furthermore, we show that *P. falciparum* is able to detect differences in host environment and adapt to it by changing the expression of genes required for host cell remodelling.

## Introduction

*Plasmodium* infection and associated mortality remain an important concern to the developing world with 218 million malaria cases and ∼450,000 deaths annually^1^ (https://www.who.int/malaria/publications/world-malaria-report-2018/en/). Widespread drug-resistance ^2-3^ and evolution of newer phenotypes; influenced by factors such as changing availability and distribution of insect vector^4^, haematological malignancies(1, 2)^5-6^ (thalassemia, sickle cell anaemia, G6PD deficiency etc), providing protective immunity to certain populations^7^, adversely impact malaria eradication campaigns. Furthermore, zoonotic infections from non-human primates, as in the case of *P. knowlesi*, is widely reported across Southeast Asia^8^ indicating a spectrum of obscured disease manifestations challenging the developing world.

*Plasmodium spp*. demonstrate an overall propensity towards immature reticulocytes for asexual development. This is evident from invasion preference of rodent parasites such as *P. berghei*, known to infect reticulocytes with ∼150-fold higher efficiency^9^. *P. falciparum*, which is responsible for the most severe form of human malaria, is able to infect red blood cells (RBCs) of all ages, yet with a preference to younger RBCs and reticulocytes ^10^. In contrast, *P. vivax* is restricted to a sub-population of reticulocytes marked by surface transferrin receptor^11^ (CD71) and are unable to infect mature RBCs. *P. knowlesi*, while able to infect all stages of RBCs in their natural macaque hosts, switch their invasion preferences to human reticulocytes during *in vitro* adaptation^12^. Since erythrocytic development is the rate limiting step in defining parasite density, transmission and disease outcome, the contribution of reticulocyte infection in parasitic behaviour and adaptation remain to be investigated.

*Plasmodium spp*. have simplified metabolic capacity since they are auxotrophic for purines^13^, vitamins and many amino acids. However, key pathways such as glycolysis, tricarboxylic acid cycle (TCA), lipid synthesis, pentose phosphate pathway, pyrimidine biosynthesis and glycosylation are conserved in these organisms^14-15^. To acquire nutrients, parasites establish new permeation pathways in the host cells. Furthermore, *Plasmodium* parasites remodel the host cells to avoid immune and mechanical clearance. Many intriguing aspects of parasite-host interactions are well studied *in vitro* in the case of *P. falciparum* using mature RBCs as host cells.

Reticulocytes have vastly different biochemical composition and properties. They contain organelles which are expelled during maturation, through exocytosis, autophagy and rearrangement of cytoskeleton^16-18^. Reticulocytes possess mitochondria with a complete complement of enzymes, including an active TCA cycle^19^ and are able to utilize glucose through the anaerobic Embden-Meyerhof pathway and hexose monophosphate shunt(3, 4). Furthermore, remnants of the transcriptional and translational machinery are also present in reticulocytes. On the contrary, normocytes retain metabolic processes needed for cellular survival -such as glucose oxidation and ion mobilization across electrochemical gradients to maintain native hemoglobin conformation^21^. In this context, the impact of two significantly different^22^ host cell microenvironments: that of immature reticulocytes and mature normocytes, on *P. falciparum* development remains an unexplored, yet critical aspect of parasite biology.

Using reticulocytes and normocytes as *in vitro* host cells, we performed a comparative study on invasion, proliferation, drug sensitivity and host-dependent adaptations of *P. falciparum*. Our results highlight the significant differences in sensitivity of parasites invaded into reticulocytes, for antimalarials artemisinin and dihydroartemisinin. Through a 100-cell transcriptomics study, we report key differences in the gene expression profiles associated with metabolism, antigenic variation and host cell remodeling in parasites replicated in reticulocytes. These results form an important dataset on which further investigations could be developed pertaining to the range and dynamics of parasite-host interactions with implications in progressive drug resistance and zoonosis.

## Results

### CD71^+^ reticulocytes support higher invasion of *P. falciparum*

We used umbilical cord blood (KK Women’s and Children’s Hospital, Singapore) for this work, since peripheral blood is not an ideal source for reticulocytes in sufficient quantities. Using a magnetic selection procedure leveraging on surface expression of CD71, we purified young reticulocytes (CD71^+^, magnet bound-fraction) and normocytes (CD71^-^, flow through fraction). Through sub-vital staining (**Fig. 1A**) and differential interference contrast (DIC) imaging (**Fig. 1B**), separation of the red cell sub-populations was confirmed, with an estimated 90% purity as reported in prior work^22^. Through western blotting (**Fig. 1C**) and immunofluorescence microscopy (**Fig. 1D**), robust separation of the sub-populations was validated.

**Figure 1.**
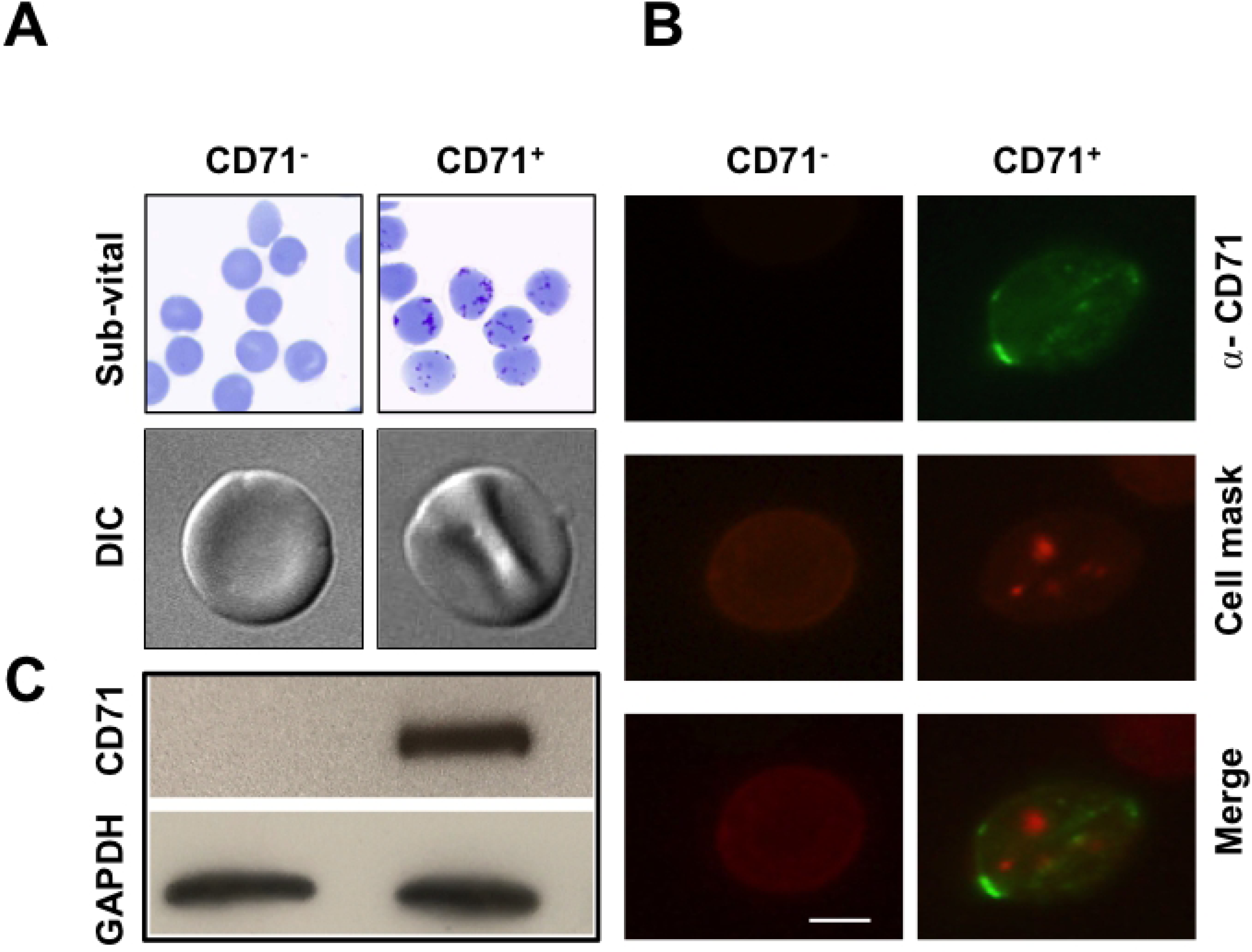
Purification and characterisation of CD71^+^ and CD71^-^ red blood cells. **A.** CD71^+^ reticulocytes were purified from cord blood using Magnetic-activated cell sorting (MACS) protocol. Isolated sub-fractions were inspected through sub vital staining (Top) and differential interference microscopy (Bottom). **B.** Immunofluorescence microscopy confirmed abundant localisation of CD71 on immature reticulocytes (magnet-bound fraction). Cells were stained with Cell Mask™ Red (Thermo Fisher Scientific) and α-CD71 antibody (Green). **C.** Western blotting showed robust purification of CD71^+^ reticulocytes. Probing was performed with α-CD71 antibody (1:1000, Abcam), with GAPDH (1:1000, Abcam) serving as loading control.

We estimated the comparative invasion efficiency of *P. falciparum* into CD71^+^ and CD71^-^ host cells. Magnet-purified schizonts (∼ 40 hpi) were introduced to both host cell types and parasitemia was determined 25 h later by counting ring-stage infections. A ∼2-fold higher parasitemia (**Fig. 2A**) in CD71^+^ cells was observed. However, the number of daughter merozoites formed were comparable irrespective of the host cell (**Fig. 2B**). Interestingly, when schizonts from CD71^+^ cells were isolated and allowed to invade CD71^-^ cells, invasion rates similar to the controls was observed, further indicating the normal rate of parasite multiplication in CD71^+^ cells. Gradual CD71 loss was recorded over 48 h (**Supplementary Fig. S1A)** for healthy reticulocytes. Upon infection, CD71 expression was mostly unchanged up to ∼18 hpi (**Supplementary Fig. S1B and S1C)** in contrast to rapid (∼ 3 hpi) maturation in *P. vivax* infection^11^.

**Figure 2.**
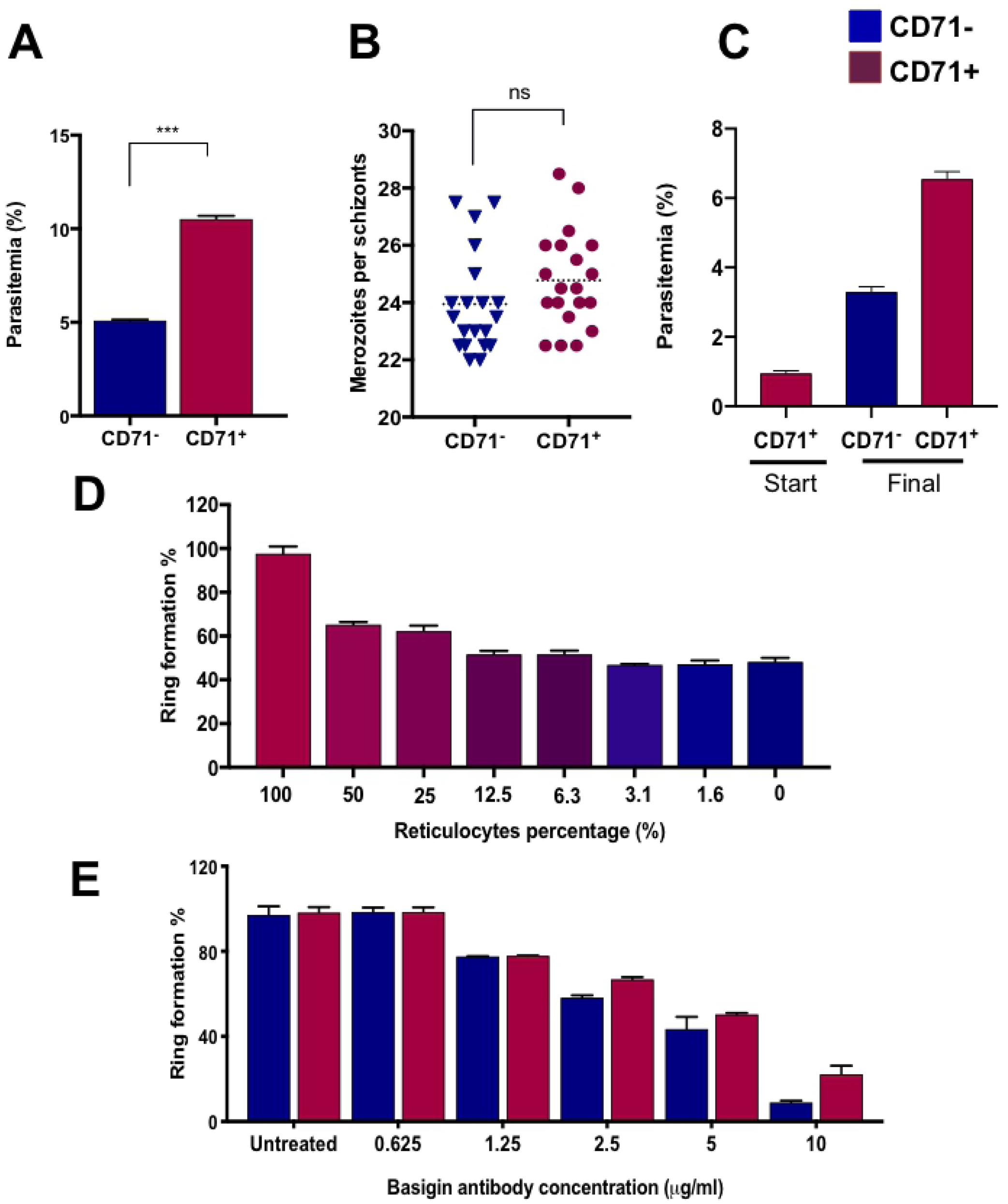
Comparison of *P. falciparum* infection in CD71^+^ and CD71^-^ blood cells. **A.** Approximately 2-fold higher infection of *P. falciparum* was observed when CD71^+^ reticulocytes (purple) were used as host cells. (Mean values of 2 independent experiments; CD71^+^: 10.52% and CD71^-^: 5.08%). An unpaired two-tailed t-test was performed to confirm statistical reliability. **B.** Merozoite counts were taken from late stage segmented schizonts through fluorescence microscopy, which showed comparable values irrespective of the host cells used for infection (Mean values of 2 independent experiments; CD71^-^: 24.28 and CD71^+^: 24.80). **C.** Parasites grown in CD71+ reticulocytes for one cycle were re-introduced to CD71-cells, which showed invasion rates similar to controls, suggesting merozoite production is not increased during development inside reticulocytes. **D.** Higher amounts of reticulocytes in the culture wells resulted in increased infection rate *in vitro*. However, maximum infection was achieved only in a pure CD71^+^ reticulocyte population. **D.** Invasion efficiency comparison in presence of varying amounts of anti-basigin antibody, saturating levels of the antibody showed differential effects on reticulocyte invasion while completely blocking *P. falciparum* invasion into normocytes.

Next, we carried out invasion assays in presence of varying ratios of CD71^+^ and CD71^-^ host cells. With an initial seeding of 1% magnet-purified schizonts, 4% rings in 100% CD71^-^ cells and 7.5% in 100% CD71^+^ cells were recorded. As expected, higher invasion correlated with higher amount of reticulocytes (**Fig. 2D**), with maximum infection in 100% CD71^+^ cells. These results suggest that despite a clear preference for reticulocytes, the selection is not entirely an active parasite-driven process but likely depends on the availability and proximity of CD71^+^ host cells. Furthermore, we did not observe higher incidents of multiply infection in reticulocytes.

Engagement of host receptors is a key step during plasmodium invasion^35^. Prior work from our group has profiled human reticulocyte proteome which indicated marginally higher amounts of basigin on reticulocyte surface (∼ 1.28-fold in comparison to mature RBCs)^22^ (**Supplemental Fig. S2**). As anti-basigin antibodies were shown to inhibit *P. falciparum* invasion^36^ we performed invasion inhibitory studies which revealed comparable invasion of both host cells at low antibody concentrations (**Fig. 2E**). However, at higher antibody concentrations (5-10 μg/ml reported from literature), invasion into CD71^+^ cells was less affected. This could be due to a combination of marginally higher basigin expression and larger surface area for interactions on CD71^+^ reticulocytes.

Hence, morphology and properties of CD71^+^ and CD71^-^ cells were determined using single cell holotomographic analysis^25^. These analysis confirmed distinct invaginations at various planes (indicated with white arrows) (**Fig. 3A**), differentiating both cell types. Evidently, CD71^+^ cells were irregular in shape with higher Refractive Index (RI), indicative of a denser cytoplasmic composition (**Fig. 3B**) consistent with previous data by Park and colleagues^25^. Furthermore, CD71^+^ cells were significantly larger (∼21%) than normocytes (**Fig. 3C**), thereby presenting larger surface area for the distribution of receptors presumably leading to easier detection and attachment by plasmodial merozoites.

**Figure 3.**
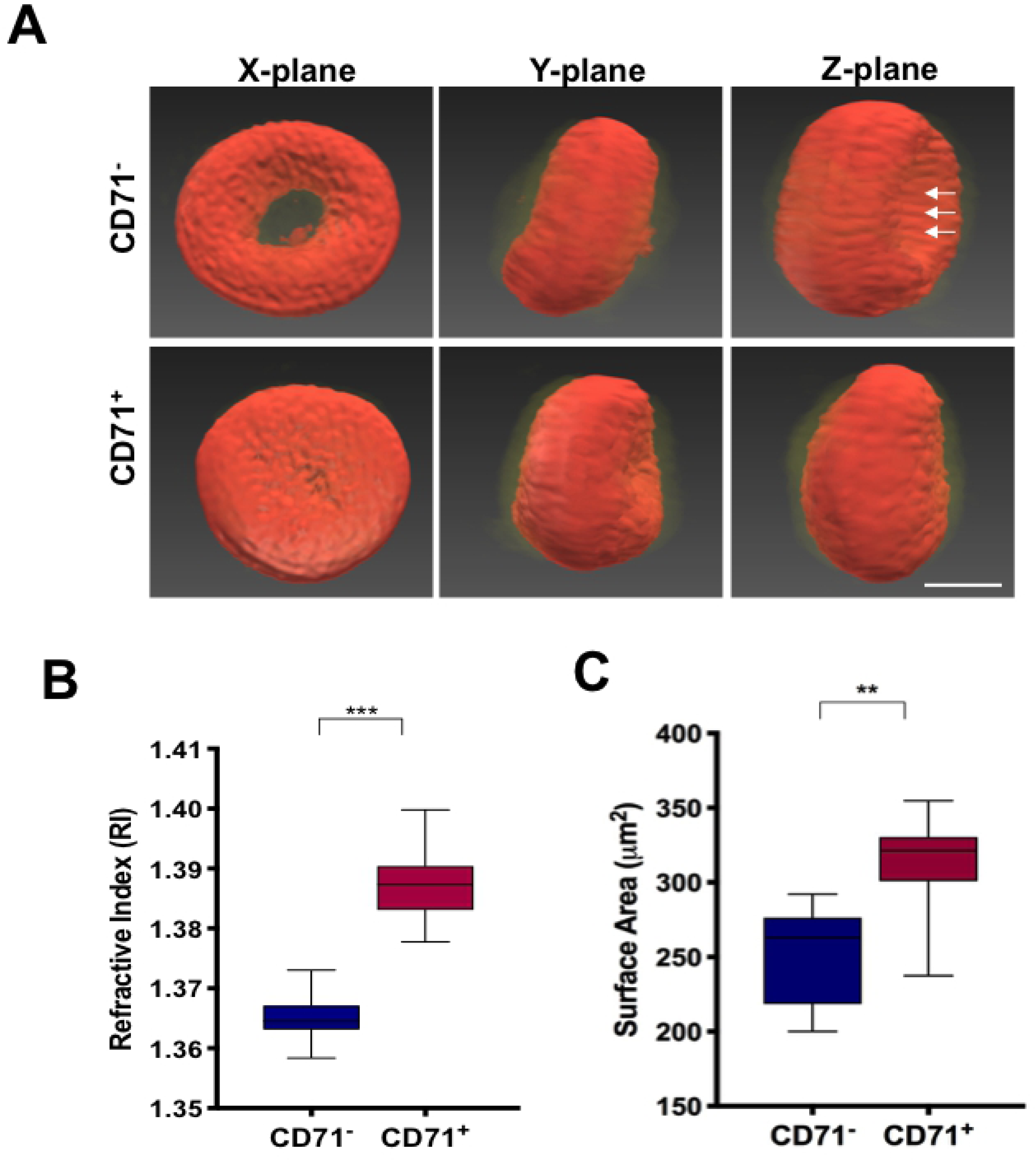
Holotomography measurements demonstrate that CD71^+^ reticulocytes are larger and multi-lobular. **A.** Representative holotomography images of CD71^+^ and CD71^-^ cells demonstrating comparative cellular morphologies. Using Tomocube HT-1, we derived quantitative dynamic cell measurements on **B**. Refractive index (Mean values of 20 cells; CD71^-^: 1.37 and CD71^+^: 1.40) and **C**. surface area (Mean values of 20 cells; CD71^-^: 238.00μm^2^ and CD71^+^: 267.75μm^2^). These measurements revealed that CD71^+^ reticulocytes are ∼20% larger than normocytes.

### *P. falciparum* grown in reticulocytes show distinct drug sensitivity profiles

Reticulocytes possess complex membranous composition^37^ and architecture rendering increased rigidity thereby influencing membrane permeability and molecular transport^38^. Previous studies show that reticulocytes have increased cation permeability for calcium (43-fold) and sodium (6-fold)^39^. Furthermore, the increased metabolic activity of CD71^+^ reticulocytes may influence metabolism of molecules, including antimalarial drugs. In this context, we selected a broad spectrum of antimalarials (along with E64 and heparin^40^, inhibitors of egress and invasion respectively) and estimated their inhibitory potential against parasites cultured in CD71^+^ and CD71^-^ host cells.

Trophozoite stage parasites (24-26 hpi) (or schizonts at 40-42 hpi for E64 and heparin) grown in CD71^+^ and CD71^-^ host cells were incubated with the drugs, along with non-treated infected RBCs. New infections were counted in the next cycle (after 50-52 h post drug treatment) and/or post invasion (20 h after treatment), for heparin and E64, followed by IC_50_ determination^26^. We observed comparable response and IC_50_ values between parasites infected into CD71^+^ and CD71^-^ cells for most drugs (**Supplemental Table 1**) including chloroquine. Interestingly, heparin blocked parasite invasion into CD71^+^ cells more efficiently (IC_50_ of 1.51 μg/ml) compared to normocytes (IC_50_ of 2.82 μg/ml), while E64 showed similar egress inhibition. In contrast, cycloheximide demonstrated remarkable killing ability against parasites in CD71^+^ cells with an estimated IC_50_ of 8.14 while infected normocytes showed an IC_50_ of 37.9 ng/ml. Host cell environment appeared to impact sensitivity of *P. falciparum* to ART and DHA (**Fig. 4A-B**), as CD71^+^ iRBCs coped with the drug exposure better. These changes in drug sensitivity is not influenced by the different invasion rates to host cells, as confirmed by tracking parasite response to drugs during stage transition (**Supplemental Fig. S3**). Although severe malaria in children is shown to cause reduced erythropoietic responses^41^, reticulocyte compensation following anemia together with sub-lethal exposure of artemisinin of parasites within CD71^+^ reticulocytes, may render progressive drug resistance.

**Figure 4.**
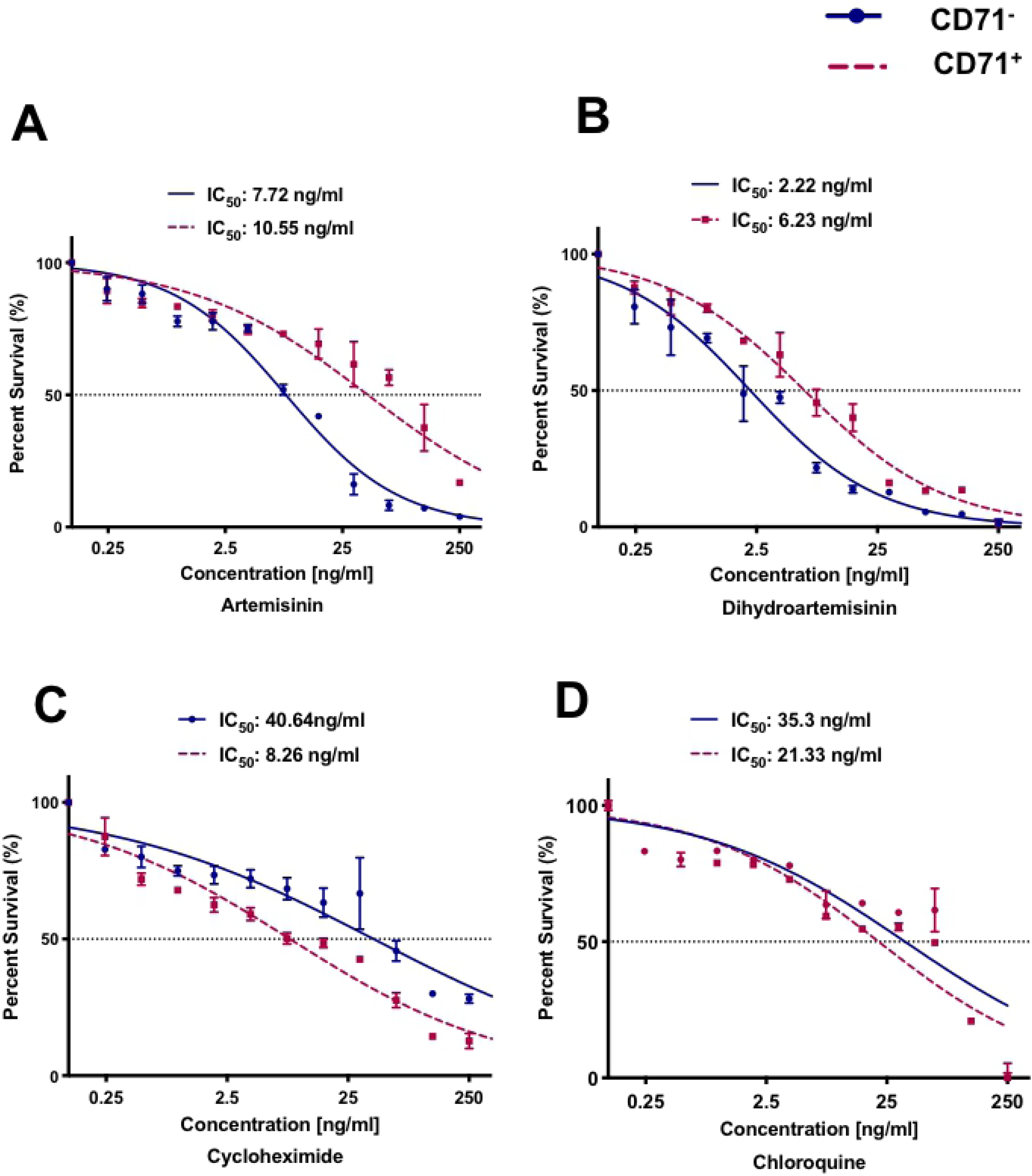
*P. falciparum* grown in CD71^+^ retioculocytes show different sensitivity to artemisinin family drugs. Growth inhibition assays against 3D7 parasites infected into distinct host cell populations was conducted. Infected cells were incubated with antimalarials at the early trophozoite stage (32 to 36 hpi) and parasitemia was manually counted (5000 cells) in the next cycle of trophozoite stage. The experimental data represents the mean of 3 independent experiments performed in replicates. Figure shows the log (inhibitor)-versus-response curves for potent antimalarials on *P. falciparum* infected CD71^+^ and CD71^-^ cells for **A**. artemisinin, **B.** dihydroartemisinin, **C.** cycloheximide and **D.** chloroquine.

### Host-dependent gene expression profiles in *P. falciparum*

*Plasmodium spp.* rely on host RBCs for metabolic needs through the consumption of host hemoglobin, vitamins and other intermediates for synthetic pathways and energy metabolism^42-44^. In addition, a range of host cell remodeling events leading to alterations in antigen presentation^45^, deformability^46^ and cytoadhesive properties^47^ are essential for the parasite to survive within the human host. Furthermore, parasites also co-opt host cell proteins to protect from damage, stage transition and proliferation^(6, 7)-49^. Hence, we set out to investigate how the different host cell microenvironments impact parasite behavior. Owing to scarcity of samples, we adopted a modified100-cell microarray technique^27^ to study gene expression profiles.

We introduced magnet-purified parasites into CD71^+^ and CD71^-^ cells as described in **Scheme. 1**. Trophozoites (100 infected cells) were sorted through FACS (**Fig. 6A**) for microarray. Clearly, differential gene expression in cellular pathways associated with protein and nucleotide metabolism, virulence as well as host remodeling (**Fig. 6B**) were observed. We identified 151 genes to be differentially expressed in CD71^+^ (cycle 1) samples such as up-regulation of translation initiation factors (PF3D7_1312400, PF3D7_0528200), tRNA ligases (PF3D7_1336900, PF3D7_0407200), pyrimidine metabolism: aspartate carbamoyltransferase (PF3D7_1344800) and orotate phosphoribosyltransferase (PF3D7_0512700). On the contrary, genes such as pantothenate transporter (PF3D7_0206200) and cysteine desulfuration protein SufE^50^ (PF3D7_0206100) involved in acetyl coenzyme A formation were significantly down-regulated in parasites infected to CD71^+^ cells (**Fig. 6C**).

**Figure 5:**
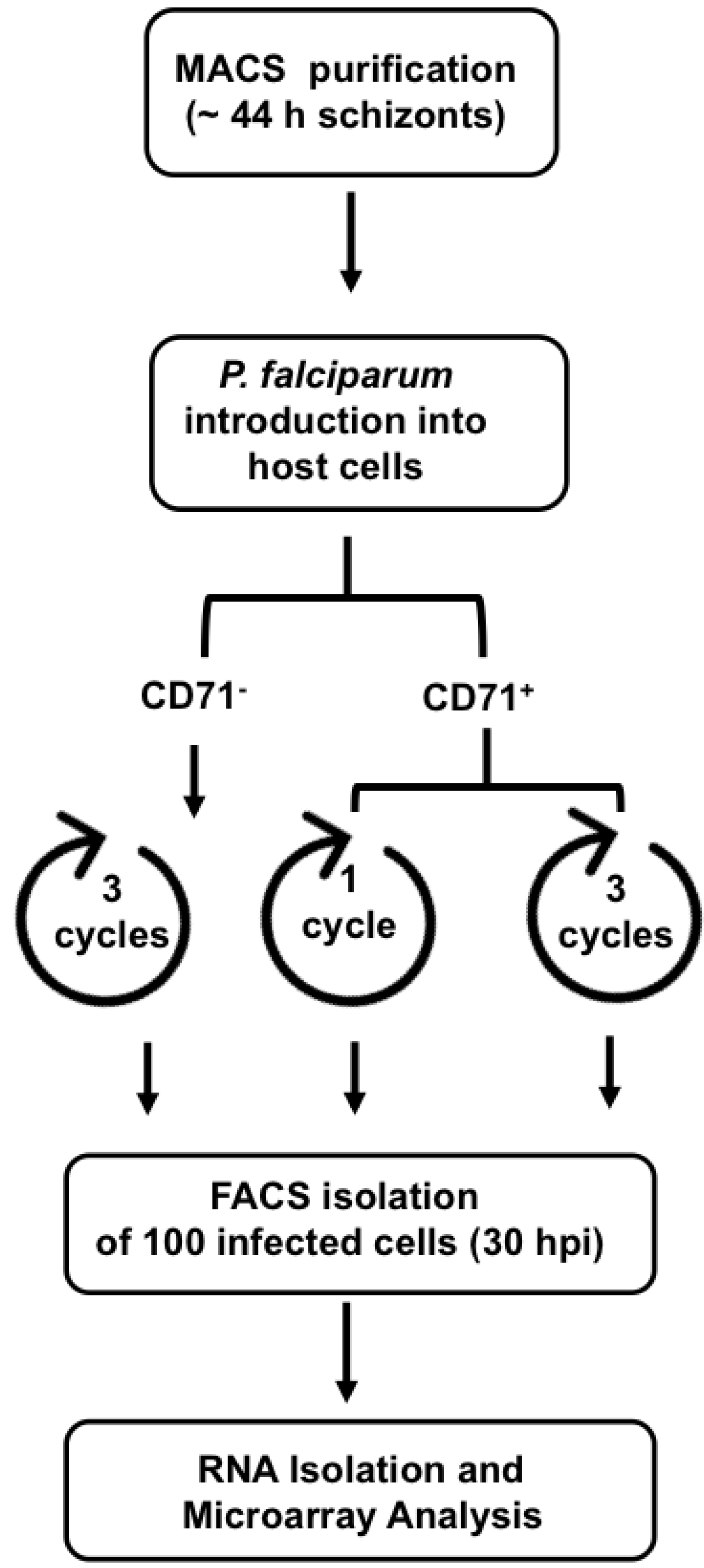
Schematic representation of parasite adaptation and enrichment strategy adopted for 100- cell microarray analysis. *P. falciparum* schizonts were allowed to invade into cord blood normocytes (for 3 continuous cycles), CD71^+^ reticulocytes (for 1 cycle) and in CD71^+^ reticulocytes for 3 consecutive cycles. For the third condition, schizonts in every cycle was purified and re-introduced into freshly purified reticulocytes (to avoid any invasion to normocytes resulting from maturation while in culture) from the same batch of blood. After staining with Hoechst, 100-infected cells were isolated by FACS sorting and subsequent microarray analysis.

**Figure 6:**
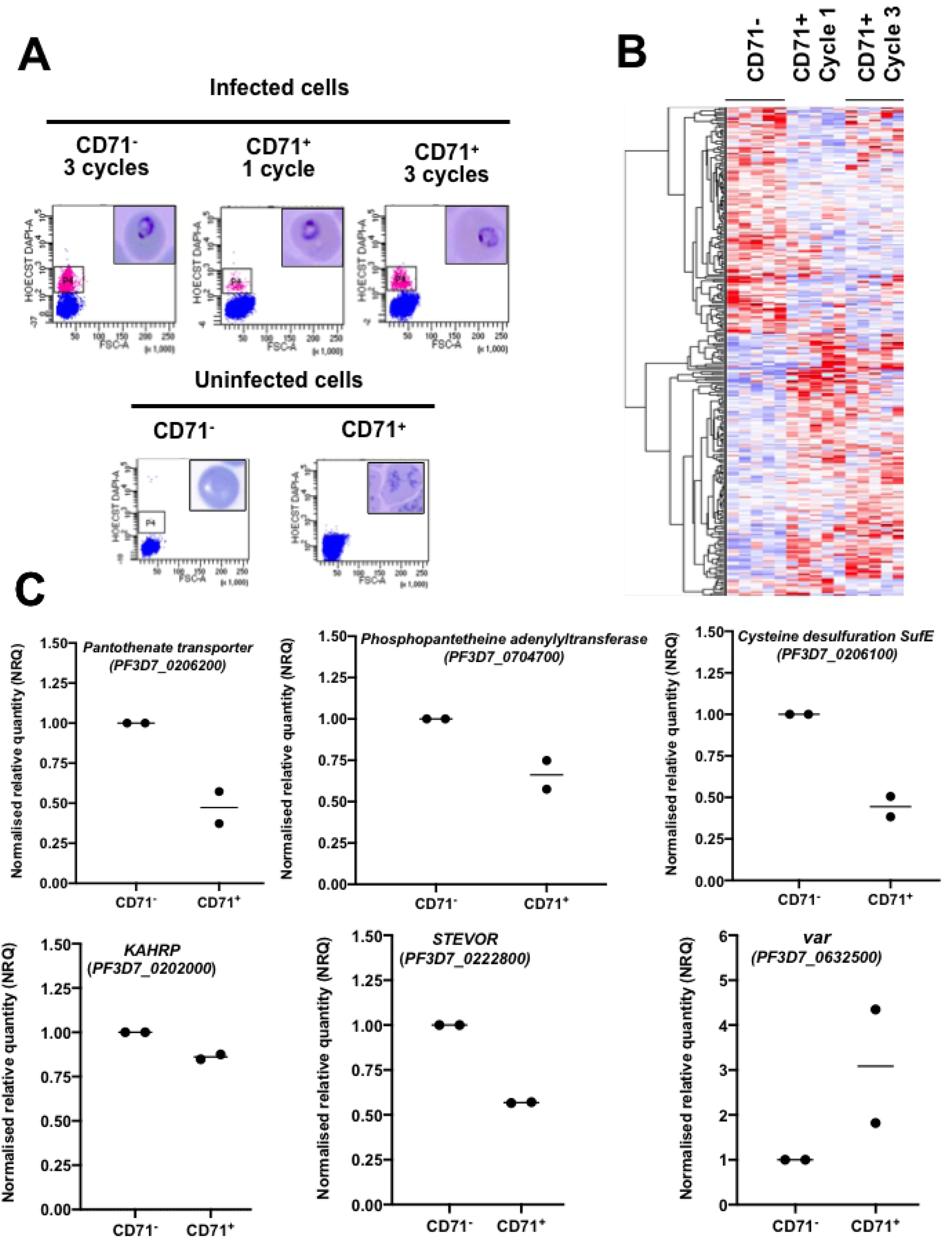
Differential Gene expression profiles in *P. falciparum* adapted in human reticulocytes. Expression profiles of genes in *P. falciparum* adapted into CD71^+^ and CD71^-^ cells. **A.** 100 infected cells from CD71^-^ cultures (3 cycles), and CD71^+^ cultures (1 cycle and 3 cycle) were isolated through FACS sorting, flow plots demonstrating gating and purity of cells (top panel). Uninfected/healthy cells were used as negative control for gating (bottom). **B.** Upon completing microarray analysis, a Z-score cut-off of 2 was applied to only assess genes, which showed stable difference between the two conditions when comparing both biological replicates. List of genes which are more than 2-fold up or down-regulated when comparing CD71^-^ host cells to CD71^+^ reticulocytes cycle 1 or cycle 3 was obtained. This analysis gave 156 genes different between normocytes versus reticulocytes in 1 cycle and 159 genes different between CD71^-^ versus CD71^+^ in 3 cycles represented in the above heatmap (based on Euclidean distance) shows the expression for these differentially expressed genes in the three conditions compared. Red represents up-regulated genes, purple represents down-regulated genes and white represents no change in expression. (CD71^-^ refers to normocytes, 3 cycles, 5 replicates; R1: refers to CD71^+^, 1 cycle, 5 replicates; R3 refers to CD71^+^, 3 cycles, 5 replicates). **C**. Based on microarray data, the differential expression of six selected genes were confirm by quantitative real-time PCR: *KAHRP* (PF3D7_0202000), *stevor* (PF3D7_0222800), *var* (PF3D7_0632500), *pantothenate transporter* (PF3D7_0206200), *phosphopantetheine adenylyltransferase* (PF3D7_0704700) and cysteine desulfuration SufE (PF3D7_0206100) confirming microarray results. Normalised quantitative expression to *arginyl-tRNA synthetase* was compared between infected CD71^-^ (control) and infected CD71^+^ (treatment). The graphs show two data points from two independent biological experiments.

3 continuous cycles of adaptation in CD71^+^ cells led to differential expression of 26S proteasome regulatory subunits (PF3D7_1306400, PF3D7_0312300), transcription regulation and mRNA splicing related genes, such as, putative DNA-directed RNA polymerase II (PF3D7_1304900), RNA topoisomerase III (PF3D7_1347100), transcriptional regulatory protein sir2b (PF3D7_1451400) and small nuclear ribonucleoprotein-associated protein B, putative (PF3D7_1414800). Additionally, genes involved in purine and pyrimidine metabolism, such as, ribonucleoside-diphosphate reductase small chain, putative (PF3D7_1015800), adenosine deaminase (PF3D7_1029600), aspartate carbamoyltransferase (PF3D7_1344800) and orotate phosphoribosyltransferase (PF3D7_0512700) were also upregulated in cycle 3 CD71^+^ parasites.

Several genes linked to host cell interaction and remodeling were differentially regulated in parasites inside CD71^+^ reticulocytes. A member of the *stevor* multigene family (PF3D7_0222800), responsible for host invasion and rosetting^51^ showed reduced expression in CD71^+^ reticulocytes. In addition, *P. falciparum* erythrocyte membrane protein 1 (*Pf*EMP1) of the *var* family showed distinct patterns with PF3D7_1255200 showing reduced expression while PF3D7_0632500 showing up-regulation in CD71^+^ host cells. Similarly, expression of knob-associated histidine-rich protein (KAHRP) (PF3D7_0202000), a major component of adhesive knobs^52^ on iRBC surface was slightly down regulated in CD71^+^ host cells, further validated through qPCR analysis (**Fig. 6C**), suggesting altered host remodeling in parasite-infected CD71^+^ reticulocytes.

### Altered host cell remodeling in *P. falciparum* infected reticulocytes

From microarray results, further validated through qPCR analyses, it appeared that expression of STEVOR, *Pf*EMP1 and KAHRP were differentially regulated which can contribute to antigenic, deformability, and cytoadherence properties of the infected cell. This was particularly interesting since we have shown in prior work that reticulocyte membrane is significantly rigid with different composition and assembly patterns of cytoskeletal components(5). Furthermore, our results also showed that infected CD71^+^ cells maintain their characteristics up to ∼20 hpi, with no indications of rapid maturation reported in the case of *P. vivax*. To elucidate possible correlations, we measured membrane deformability properties of *P. falciparum* infected CD71^+^ and CD71^-^ cells. Membrane of the healthy CD71^+^ reticulocytes were significantly rigid with an estimated membrane shear modulus of 19 pN/μm compared to normocytes (average smear modulus of 6.5 pN/μm), as reported^22-37^. *P. falciparum-*infected CD71^+^ cells remained rigid during the progression of parasites (**Fig. 7A**) in agreement with CD71^+^ signal for up to ∼20 hpi (**Supplemental S1B-C**). These findings support that *P. falciparum* is able to sense a stiffer host cell membrane (CD71^+^) and calibrate remodeling events by altering gene expression, particularly for members of the STEVOR and *Pf*EMP1 family that are involved in these processes.

**Figure 7:**
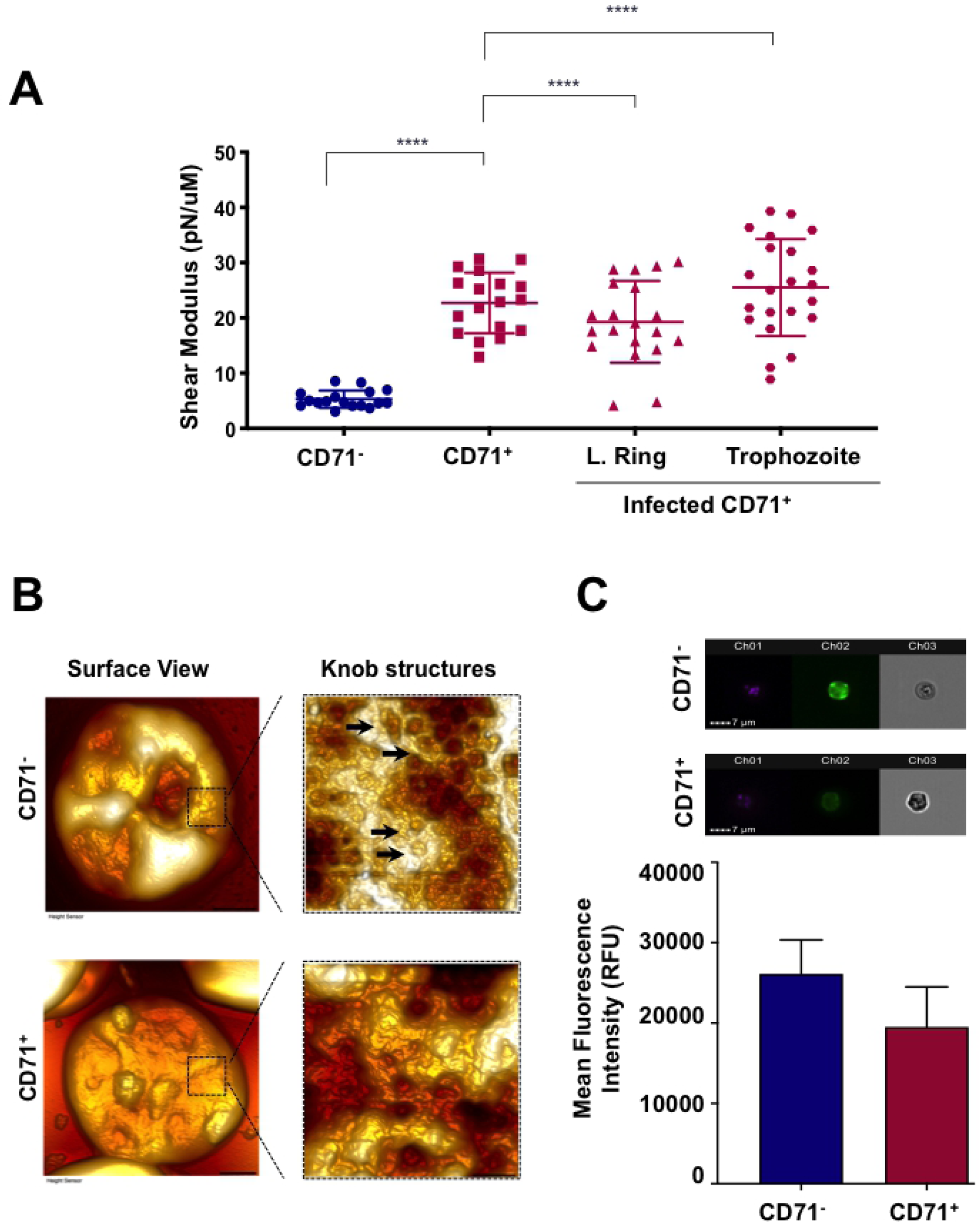
Host cell remodelling in *P. falciparum* infected reticulocytes is altered through differential gene expression. **A.** Membrane of *P. falciparum* infected reticulocytes (CD71^+^) remain stiff during early parasite development, as measured through micropipette aspiration technique. Briefly, 30-40 cells were obtained per measurement for individual experiments. The cellular membrane was monitored by Olympus IX71 microscope and image was processed by QCapture Pro 6.0 and the elastic shear modulus was determined using the Hochmuth model. **B.** Surface view of infected CD71^-^ and infected CD71^+^ cells scanned by AFM showing an area of 1µm on a representative cell, indicating reduced knob formation in infected CD71^+^ reticulocytes.**C**. Reduced KAHRP expression on infected CD71^+^ cells was validated through Amnis Imaging flow cytometry. Infected cells were labeled with anti-KAHRP antibody and stained with secondary anti-mouse FITC antibody together with Hoechst. Background FITC fluorescence was deducted from doubly-stained uninfected healthy cells: CD71- and CD71+, as appropriate. Graphs shows results from three independent experiments.

Gene profiles also suggested down-regulation of KAHRP expression, which by forming cell surface knobs facilitates endothelial adhesion. In general agreement with the data, the cell surface images generated through Atomic Force Microscopy (AFM) indicated reduced distribution of knob structures (**Fig. 7B**) on the surface of infected CD71^+^ reticulocytes. To further quantify these differences, we estimated KAHRP expression on infected CD71^+^ and CD71^-^ cells through imaging flow cytometry using an antibody against KAHRP, which confirmed reduced signal from CD71^+^ cells infected with *P. falciparum.* We were able to measure ∼20% reduction in KAHRP expression in infected CD71^+^ cells (**Fig. 7C**).

## Discussion

*P. falciparum* and *P. vivax* present contrasting cases in terms of distribution, disease severity^53^, drug sensitivity/resistance^54^ and relapse^55^. Perhaps, the most intriguing aspect that differentiates these two species is the strict restriction of *P. vivax* to immature reticulocytes^11^. Since reticulocytes are formed in the bone marrow and only found in small numbers in the peripheral circulation, higher bone marrow parasitemia were reported for *P. vivax*, despite blood examination showing no infection^56^. Within the bone marrow, these parasites remain undetected and mature rapidly into transmissive forms^57^. Interestingly, homing of *P. falciparum* gametocytes in bone marrow has also been a topic of recent investigations^58^. Positive correlation between anaemia and bone marrow hemozoin/parasites in children with severe malaria reported by Aguilar *et al*^*59*^ and others^60^, indicates that bone marrow reticulocytes remain susceptible to *P. falciparum* asexual stage development.

Reticulocyte infection is advantageous for *P. falciparum* due to the abundant resources for energy metabolism and likely protection from chemical/oxidative damage. While our results do not indicate increase in *P. falciparum* multiplication rate in reticulocytes, but they appeared to better cope with damaging effects of drugs *in vitro*. Although, apparent differences in the membrane permeability and likely metabolic fate for drugs in the reticulocyte cytoplasm cannot be ruled out as a contributing factor. While RBC damage and resultant anaemia ensues over production of reticulocytes, it offers *P. falciparum* an opportunity to invade reticulocytes. Reticulocytes contain higher amounts of hemoglobin ^25,61^, which the parasites catabolize generating hemozoin. Artemisinin and combinations thereof target hemozoin pathway and thus required in higher dozes against *P. falciparum* residing within reticulocytes. In the light of our results, contribution of host reticulocytes in progressive drug resistance warrants systematic dissection.

Energy metabolism in *Plasmodium* depends on resources available through host RBCs and beyond. For example, pantothenic acid, an essential precursor for Coenzyme A is scavenged from host RBCs or culture media^62^, uptake of which is facilitated by pantothenate transporter (PF3D7_0206200). We observed reduction in the expression of this gene in *P. falciparum* infected reticulocytes. Furthermore, gene coding cysteine desulfuration SufE (PF3D7_0206100), which is also linked to Coenzyme A biosynthesis was down-regulated in these parasites. This is interesting since the parasites invaded into reticulocytes have an overall nutrient/energy rich environment and may not be required to import and utilize extracellular pantothenate. While pantothenate transporter is refractory to deletion in *P. falciparum*, its role is primarily assigned to sexual differentiation^63^ and transmission. Down-regulation of pantothenate transporter and associated proteins (for example phosphopantetheine adenylyltransferase) may indicate lesser commitment for sexual stage transition having sensed a rich environment for asexual proliferation. However, this hypothesis remains to be further developed.

Cytoskeletal rearrangements leading to actin remodelling through exported proteins such as *Pf*EMP1, RESA^64^ and STEVOR^65^ is a hallmark of *P. falciparum* asexual development. KAHRP plays a critical role in iRBC rigidification through the formation of knobs^66^, accounting for 50% of the increased shear modulus. These changes are essential for parasites that are in the peripheral circulation to avoid both immune and mechanical clearance. Although *in vitro* culture conditions are known to affect transcriptional profiles^67^, our observations that several members of these gene families are differentially regulated implies that the parasites are able to detect a vastly different host cell properties, and adapt to it by changing gene expression. Afterall, if the primary location of infection remains the bone marrow where reticulocytes are abundant, parasites are not threatened by splenic clearance. Furthermore, *P. vivax* restricted to reticulocytes do not form knobs^68^ and completely lack homologs of *var* genes, offers additional premises to undertake these observations for future research. Evolution of new biological tools such as the ability to engineer and cultivate homogeneous populations of young red blood cells for functional dissection of host factors^69-70^ critical for parasite invasion, development and adaptations may contribute to such efforts.

## Materials and Methods

### Ethics Statement

All plasmodium experiments were conducted with approved protocols from the Institutional Biosafety Committee (IBC) of the Singapore University of Technology and Design (SUTD). Blood for routine parasite culturing and maintenance was purchased from Interstate blood bank, USA. Cord blood samples from adult normal term pregnancies were collected at KK Women’s and Children’s hospital with written informed consent. Protocol to collect and use cord blood for experiments was approved by the SingHealth centralised institutional Review Board (CIRB). All cord blood samples were anonymized.

### Parasite culturing, synchronization and analysis

Blood was centrifuged at 600 × g for 10 min to remove buffy coat and stored in Malaria Culture Medium (MCM). Washed cord blood was incubated with CD71 magnetic microbeads for isolation of reticulocytes (CD71^+^) through MACS (Miltenyi Biotec, Singapore), as mentioned previously(5). Unbound fraction was stored at 4°C as a source of normocytes (CD71^-^).

3D7 strain of *P. falciparum* was used for all experiments. Parasites were maintained in 2.5% hematocrit in RPMI-HEPES medium at pH 7.4 supplemented with hypoxanthine 50 µg mL^−1^, NaHCO_3_ 25 mM, gentamicin 2.5 µg mL^−1^, and Albumax II (Gibco) 0.5% wt/vol. Schizonts (∼ 45 hpi) were enriched using MACS (Miltenyi Biotech, Germany) followed by selection of rings by 5% sorbitol ^23^.

Blood smears prepared on glass slides were fixed with 100% methanol (Merck) and stained with fresh 1:10 Giemsa (Merck) solution. Smears from infected CD71^+^ samples were stained with new methylene blue and examined under 100X oil immersion objective (Leica ICC50 W). Images of parasitic phenotypes were captured using a Leica digital camera^24^.

### Optical diffraction measurements using Tomocube™

Freshly collected RBC/reticulocyte samples were diluted 1:1000 with PBS/BSA (1%) for 2D Optical Diffraction Tomography (ODT) measurements. Images were acquired at multiple illumination angles using a 3D RI tomogram at an excitation at 532 nm (HT-2H, Tomocube, Inc., Daejeon, Korea), as described previously^25^ and processed using Image J.

### Antimalarial drug assays

Drugs were aliquoted at 10 mg/ml either in H_2_O (chloroquine, cycloheximide, E64 and heparin), ethanol (halofantrine) or DMSO (Dihydroartemisinin, artemisinin, atovaquone, piperaquine and trichostatin A. Schizonts (∼44 hpi) were introduced to CD71^+^ and CD71^-^ host cells separately, as seed cultures. Parasites were checked when they became trophozoites/schizonts (after ∼ 35 h) through microscopy. Schizont stage parasites (40-42hpi) from the CD71^+^ and CD71^-^ cells were isolated and mixed with appropriate fresh host cells at 1% parasitemia and 2.5% haematocrit together with drugs. Untreated infected RBCs were included as negative controls. After 40 h, parasitemia was estimated through manual counting. Three trained researchers counted the slides independently, data represents experiments performed in triplicates. IC_50_ values were determined using GraphPad Prism according to the recommended protocol for nonlinear regression of a log(inhibitor)-versus-response curve^26^. An unpaired t-test was applied to ensure statistical significance.

### Parasite adaptation and transcriptomics

Parasites grown in CD71^-^ and CD71^+^ cells (1 cycle and 3 cycles), each time purifying late stage parasites (∼35 hpi) on MACS and re-introducing into fresh host cells. In the next cycle, cells were harvested (30 hpi), stained with Hoechst for 30 min followed by washing in PBS before sorting 100 infected cells using a BD FACSAria™ (BD Biosciences, Singapore) into 0.2 ml tubes containing cell lysis buffer (RNase inhibitor and BSA). Samples were stored in -80°C. For cDNA amplification, cell lysates were subjected to SMART-seq2 protocol^27^ and cDNA was purified using QIAGEN PCR purification kit. Before microarray hybridization, 2 to 3 µg of cDNA was labelled with Cy5 (sample) and Cy3 (reference pool) dyes and incubated for 2 h. Labelled samples were purified using QIAGEN PCR purification kit and hybridized on *P. falciparum* intragenic DNA chip at 70°C for 18 h. Next day, hybridized chips were washed and scanned on Power Scanner (Tecan).

### Quantitative real-time PCR (qPCR)

qPCR was performed on two biological replicates, one set from RNA prepared for 100-cell microarray and a second fresh set of experiments. For the latter, total RNA was extracted (PureLink RNA mini kit, Life Technologies) and cleaned up through on-column digestion with PureLink DNase (Life Technologies) and reverse transcribed using the iScript™ cDNA Synthesis kit (Bio-Rad Laboratories). Real-time PCR was performed on a CFX-96 Touch System (Bio-Rad Laboratories). The PCR reactions were set up using iTaq™ Universal SYBR Green Supermix (Bio-Rad Laboratories), programmed at 30s/ 95°C followed by (10s at 95°C, 30s at 53°C) × 40. Melting curve analysis and gel electrophoresis were performed to confirm the specificity of PCR amplicons.

Gene specific primers were designed from Primer-BLAST^28^, sequences are given as **Supplemental Table-2**. *P. falciparum Arginyl-tRNA synthetase* (PF3D7_1218600) was used as reference gene^29^. We used Pfaffl method^30^ to calculate normalised relative quantity (NRQ), infected normocyte (iNorm) was considered control and infected reticulocyte was considered treatment sample. The results were from two biological replicates, each with two technical replicates.

Amplification efficiency (E) of each primer pair was calculated from E = 10^(−1/slope)^. The E of all primer pairs in this study were within 1.8 - 2.0.

ΔCt_gene of interest_ = Ct_goi_ control - Ct_goi_ treatment; ΔCt_reference_ = Ct_ref_ control - Ct_ref_ treatment.

The normalised relative quantity was calculated as below:

NRQ = (E_goi_)^ΔCt_goi_ / (E_ref_)^ΔCt_Ref_ where E_goi_ and E_ref_ are respectively the amplification efficiency of target gene and reference gene *Arginyl-tRNA synthetase*.

### Measurement of KAHRP expression on infected reticulocytes by

ImageStream Infected cells (42 hpi) were collected, washed with 1xPBS and fixed with 4% paraformaldehyde and 0.0075 % glutaraldehyde for 30 min at RT. Subsequently, cells were permeabilized in 0.1 % Triton X-100 for 5 min/RT. Following a quenching step in 0.1M Glycine for 30 min/RT, blocking was done overnight at 4° C in 3% BSA/PBS. Incubation with anti-KAHRP monoclonal antibody (mAb 18.2, European Malaria Reagent Repository) at 5 µg/ml in 3 % BSA was done for 1 h at RT. Samples were washed and incubated with 1:500 goat anti-mouse IgG FITC antibody (Abcam, ab6785) together with 1 µg/ml Hoechst 33342 (Sigma Aldrich) in 3% BSA for 1 h at RT. After washing, cells were re-suspended in 70 µL PBS and used for imaging.

Data was acquired with ImageStream X MkII flow cytometer (Merck, Darmstadt, Germany) at 60x magnification for high-content single-cell analysis^31^. Hoechst-positive (Ch 01) events were gated according to fluorescence intensity and visual inspection of images confirming only parasite-infected samples were being analyzed. From this, median fluorescence intensity (MFI) of FITC/KAHRP (Ch 02) was obtained. Background FITC fluorescence was deducted from doubly-stained healthy RBCs or reticulocytes used as blank controls. Single stain controls were prepared for compensation matrix generation by IDEAS software and applied for all 3 independent experiments.

### AFM imaging of infected RBC surface

AFM imaging was performed with Bruker Dimension FastScan microscope (Bruker) using super sharp silicon probes (SSS-NCHR probes, Nanosensor, Switzerland) in air tapping mode^32^. Height images were captured at a resolution of 512 samples per line for 1 µm × 1 µm areas with a scan rate of 0.5 – 1 Hz. NanoScope Analysis software (version 1.90) was used to generate images with sample height profiles. Images were smoothened using a low-pass filter based on Gaussian convolution kernel, resulting into topographical height images of sample surface.

### Micropipette aspiration

A micropipette with an inner diameter of 1 ± 0.25 µm was used to aspirate the RBC membrane to estimate membrane stiffness, as described in prior work^33^. A pressure drop rate of 6 Pa/s and a total pressure drop of 100 Pa were applied to aspirate and deform each cell membrane. The aspiration was visualized on a Nikon TE2000-S microscope and processed by a Labview based software. The recorded aspiration values were manually extracted, and the shear modulus was calculated using the Hochmuth model^34^.

## Acknowledgements

RN acknowledges SUTD Ph.D. Scholarship awarded by Ministry of Education (MoE), Singapore. RN, HY, TTTC, PT, GS and RC acknowledges infrastructure support through SUTD-MIT International Design Centre (IDC) and funding through T1MOE1702 and RGUOO180301 grants. Miss Faith Liew’s (KK Women’s and Children’s hospital) assistance with blood collection and Mr Benedict Lim’s (Tomocube) support with 3D RI measurements are acknowledged. JCKY received salary support from Singapore’s Ministry of Health’s National Medical Research Council (NMRC/CSA-SI-008-2016).

## Figure legends

**Supplemental Table. 1: IC**_**50**_ **values for *P. falciparum* grown under two host conditions against selected antimalarial drugs.**

**Supplemental Table. 1: A list of genes with corresponding PCR primers used for qPCR experiments**

**Supplementary Figure-1. *P. falciparum*-infected red cells remain CD71**^**+**^ **until parasites progress into late rings/early trophozoites. A.** CD71^+^ samples were stained with anti-CD71 antibody at 1:100 and secondary antibody (Alexa Fluor 488 at 1:200). Differential interference microscopy (DIC) show morphological characteristics of reticulocytes still present at ∼24 h. **B.** Fluorescence intensity of stage specific depletion of CD71 on uninfected CD71^+^ verses infected CD71^+^ reticulocytes (‘+’ indicated presence of fluorescence, ‘-’ indicated absence of fluorescence; based on 1000 infected cells). Samples were imaged on coverslips and captured at 100x oil magnification using a CKX53 Olympus microscope. **C.** Sub vital stain of cells at specific stages of infection (ring ∼18 to 24 h), trophozoite (∼24 to 36 h) and schizont (∼ 40 to 44 h) show absence of RNA (green arrow) in infected while uninfected CD71^+^ cells still harbour reticular matter (red arrows). (ui: uninfected ; i: infected). Bar graph depiction of the premature loss of CD71 on infected reticulocytes in comparison to uninfected reticulocytes.

**Supplementary Figure-2. Surface expression of known host receptors required for *P. falciparum* invasion** (Results originating from Mass spectrometry data, reported in Chu *et al*, 2018).

**Supplementary Figure-3. Differential sensitivity of antimalarial drugs between infected reticulocytes and infected normocytes is not caused by differences in the invasion rates**. Seeding cultures for both infected host cells were reduced to the same parasitemia of 1% followed by treatment of antimalarials **A.** artemisinin **B.** dihydroartemisinin **C.** cycloheximide and **D.** chloroquine at IC_50_ and IC_80_ concentrations at early trophozoite stage and parasitemia was manually counted in the next cycle of trophozoite stage (32 to 36 h post invasion).

**Supplementary Figure-4. Pathway enrichment results demonstrating altered metabolism and host cell remodelling in (A) parasites grown for 1 cycle in CD71**^**+**^ **reticulocytes and (B) parasites grown for 3 continuous cycles in reticulocytes.**

**Supplementary Figure-5: Relative mRNA levels of interseting candidate genes across 3 cycles of host cell switching (N-R1) and adpatation (R1-R3)** Genes such as **(A)** var (PF3D7_1255200), **(B)** KAHRP, **(C)** Pantothenate transporter and **(D)** orotate phosphoribosyl transferase are higlighted. Transcript levels were measure by qPCR and shown as normalised relative quantity to internal control *arginyl-tRNA synthetase* gene. Results were from one biological experiment of the RNA sample used for microarray data.

**S6. Representative images of infected CD71**^**+**^ **and CD71**^**-**^ **red cells obtained from Imaging flow cytometry, demonstrating reduced fluorescence signal in infected CD71-host cells.**

**S7. Surface patterns of healthy (un-infected) CD71**^**+**^ **and CD71**^**-**^ **red blood cells, used as controls for studying the presence of knobs.**

**S8: Genes identified and fold difference from the Microarray experiments**

## Authorship Contributions

**RN, TTTC, JT, YH, GS, JXT, PT and KF carried out laboratory work and collected and analyzed the data; JC** provided clinical management of cord blood related aspects, ethical clearance, and collection and processing of the blood samples; **RN, TTTC, JT, HY, GS, JXT, PT, KF, KST, CT, JC and ZB** participated in data interpretation and helped to draft the manuscript; **RC** designed the study, coordinated the project and wrote the manuscript. The funders had no role in study design, data collection and analysis, decision to publish, or preparation of the manuscript.

